# Dynamic Compartmentalisation of Intracellular Sodium in collecting duct cells

**DOI:** 10.1101/2022.06.20.496806

**Authors:** A. Assmus, L. Mullins, C. Sherborne, A. Peter, J. Early, F. Claeyssens, J. W. Haycock, R. Hunter, J. J. Mullins

**Affiliations:** Center for Cardiovascular Science, University of Edinburgh, Edinburgh, UK; Department of Materials Science and Engineering, University of Sheffield, Sheffield, UK; Center for Discovery Brain Sciences, University of Edinburgh, Edinburgh, UK

**Keywords:** sodium, multivesicular bodies, principal cells

## Abstract

In Principal cells (PC) of the cortical collecting duct (CCD), the highly regulated and coordinated reabsorption of sodium occurs through the epithelial sodium channel (ENaC) at the apical membrane and Na/K ATPase at the basolateral membrane. However, it is not known how sodium ions (Na^+^) are transported across the cell. We investigated intracellular transport in mCCD_cl1_ cells using a fluorescent sodium dye, CoroNa Green AM. Dye uptake was stimulated by aldosterone, blocked by amiloride (an ENaC inhibiter), and basolateral transport was prevented by ouabain (an Na/K ATPase blocker) thus validating the dye’s apparently faithful replication of sodium transport. Cells exhibited a consistent pattern of sodium-containing vesicles, of various sizes, surrounded by cytoskeleton and lipid membrane. While the smallest vesicles (∼0.5μm) co-stained with lysotracker, larger vesicles (up to 6.4μm) did not co-stain with either lysosomal- or mitochondrial-specific dyes and appeared to have internal structure, suggesting that they were multivesicular bodies. Time-lapse imaging showed a subset of these multivesicular bodies release or take up sodium dye in a controlled manner. Our novel data suggest that intracellular sodium compartmentalisation is highly regulated and offer new insights into intracellular sodium dynamics in the collecting duct, revealing potential new targets for control of sodium homeostasis.

**New and Noteworthy:** The article shows for the first time, to our knowledge, intracellular sodium transport mechanism in mCCD_cl1_ cells in the form of dynamic vesicular bodies. These structures offer new targets for the regulation of sodium homeostasis and transport in the kidney collecting duct, with wider implications for blood pressure regulation.

## Introduction

In kidney collecting ducts, the main function of principal cells is to maintain water and salt homeostasis, with high variations in luminal (urinary) sodium concentration resulting in high variations in sodium and water transported to the blood stream. At the basolateral membrane, sodium is actively extruded by Na-K ATPase - 3 Na^+^ ions cross into the blood stream but only 2 K^+^ ions enter the cell - thus generating an electrochemical gradient. Luminal sodium is transported at the apical membrane through the epithelial sodium channel (ENaC) across the electrochemical gradient. The actions of ENaC and Na-K ATPase must be co-ordinated to maintain intracellular sodium concentration within a narrow range commensurate with basic cell functions (1) whilst accommodating the variations of Na+ influx at the apical membrane. Little is known about the specific transport of sodium from apical to basolateral membrane, or its regulation, though a potential intracellular Na^+^ sensor or regulation mechanism, yet to be identified, was previously suggested (1). This gap in knowledge has implications for the general regulation of sodium reabsorption and in consequence, of blood pressure.

Due to its robust sodium and potassium transport capacity as well as response to physiological concentrations of hormones, the mCCD_cl1_ cell line is used as a model for collecting duct principal cells (2–4). In this study, we observe and investigate the appearance and disappearance of intracellular, sodium-containing, multivesicular bodies in mCCD_cl1_ cells, using an Na^+^ fluorescent dye, CoroNa Green AM, (NaG; Invitrogen) imaged in live cells. To our knowledge this is the first description of such dynamic sodium-containing structures being reported in CCD cells.

## Methods

### 3D scaffolds

Bespoke porous polymerised high internal phase emulsion scaffolds (polyHIPEs)(5) were created for culture of mCCD_cl1_ cells. To create a monolayer of cells on the scaffold’s surface, porosities were kept at a size range under 20μm, using a method described previously (6). Cylindrical scaffolds (10cm long, 1cm diameter, 1mm thick) were polymerised. To prepare for cell culture, scaffold discs were cut with a razor blade, bathed in 70% EtOH, and rehydrated in PBS, then in cell culture medium.

### Cell culture

mCCD_Cl1_ cells were cultured in phenol red-free DMEM/F-12 media (Invitrogen, Life Technologies), supplemented with: 5 μg/ml insulin, 50 nmol/l dexamethasone, 1 nmol/l triiodothyronine, 5 μg/ml apotransferrin, 60 nmol/l sodium selenite, 10 ng/ml EGF, 2% FBS, and 100 U/ml to 100 μg/ml penicillin-streptomycin (Pen-Strep) as previously described (2). As appropriate, cells were treated with aldosterone (Sigma-Aldrich A9477; 3nM for 3 hours before imaging); Amiloride (Sigma-Aldrich A7410; 10μM for 10min); Ouabain (Sigma-Aldrich O3125; 5mM for 30min). For dome formation, mCCD_cl1_ were seeded at a 1:10 split ratio in glass bottom dishes (35mm, Thermo Fisher). For culture on 3D scaffolds, cells were seeded at a 1:1 split ratio and left to attach for 3 hours before changing the culture medium. Cells were then cultured for 7 to 9 days before further experiments.

Cells were switched to a basic medium of DMEM/F-12 supplemented with Pen-Strep 2 days before aldosterone experiment.

### Tubule isolation

Kidneys from WT mice were harvested, and the cortical region cut and finely minced and incubated twice for 30min at 37°C with frequent (every 5min) shaking in a Collagenase IV solution (Thermo Fisher, 17104019. 250μg/mL in HBSS, 1.1ml/kidney). Following sedimentation of debris, the supernatant containing tubules was quenched with BSA (10mg/mL in HBSS) and transferred to cell culture medium for further experiments.

### Dyes

Live cells were stained with fluorescent dyes: CoroNa Green AM (NaG, 5μM, CoroNa™ Green cell permeant, Invitrogen C36676; 30min incubation before imaging); Potassium green (KG, 10μM, ION Potassium-Green-2 AM, K^+^ indicator, abcam ab142806; 45min incubation time), a lipophilic styryl dye (5μg/mL, FM™ 4-64, Molecular Probes T13320) for plasma membrane and vesicle formation; a polymerized tubulin dye (1μM, Tubulin Tracker™ Deep Red, Thermo Fisher Scientific T34077) for cytoskeleton staining. Cell nuclei were stained with Hoeschst 33342.

Lysosomes and mitochondria were stained with Lysotracker™ Red and Mitotracker™ Red respectively (75nM, ThermoFisher, L7528 and M22425).

### Immunolabelling of cells on scaffold

Cells cultured on polyHIPEs scaffolds disks were fixed using 4% PFA for 20 minutes. Double immunostaining *of* α-ENaC and acetylated-α-tubulin was made using a rabbit anti-mouse α-ENaC antibody (1:1000, kindly provided by J.Loffing, University of Zurich, Switzerland) with Alexa Fluor 568 secondary antibody (1:500), and an anti-acetylated-α-tubulin conjugated antibody (1:50, Santa-Cruz Biotechnology sc-23950 with AF488, green secondary).

### Imaging

For doming cells and isolated tubules, we used a Leica TCS SP8 confocal microscope, with the ×10, x20 (Air, NA 0.40 and 0.75 respectively) and x63 (Oil, NA 1.4) objectives. The 405, 488, 594nm lasers were used for detection of the relevant dyes described above. For cells cultured on PolyHIPE scaffolds and time-lapses, we used a Zeiss LSM880 microscope with Airyscan unit. The x63 (Oil, NA 1.4) objective was used, with the 405 and 488 lasers for detection of Hoechst and NaG.

### Image analysis

Images were analysed using ImageJ (Institute of Health) or Imaris Imaging Software (Oxford Instruments).

## Results

When mCCD_cl1_ cells are cultured on glass-bottom dishes (or in flasks or plates), they form ‘domes’. After reaching confluency a subset of cells polarise, transport water and solutes, and detach from the substrate, forming a monolayer of cells around an increasing volume of filtrate (See Fig. 1A). Application of NaG showed that Na^+^, together with NaG, is concentrated in these domes as expected (Fig. 1B). Closer inspection of the cells on or around the domes with confocal microscopy showed intracellular “droplets” or vesicles containing NaG.

**Figure 1:**
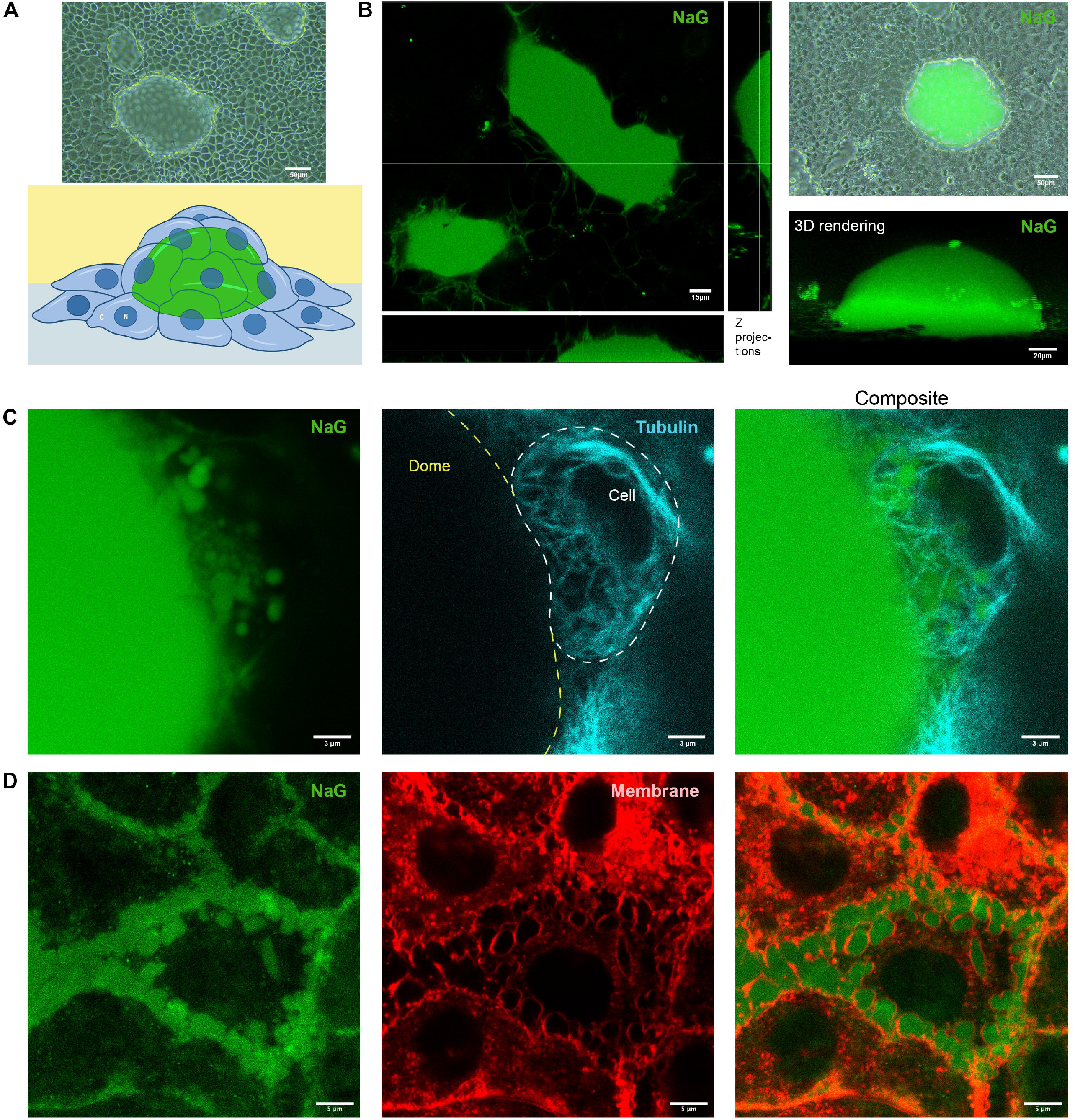
A. Top panel: brightfield image of mCCD_cl1_ cell culture. Lighter, round shapes correspond to ‘domes’ of cells detached from the plate. The doming monolayer encloses green filtrate as shown in the cartoon below. B. Left panel: z-projection rendering of NaG fluorescence in doming mCCD_cl1_ cells. Top right: overlay of brightfield and green channels of NaG in doming mCCD_cl1_ cells. Bottom right: Reconstituted 3D view of dome. C. Image of live mCCD_cl1_ cells stained with NaG (green) and Tubulin dye (light blue) on the side of a dome after aldosterone treatment. D. Staining of live mCCD_cl1_ cells using NaG (green) and lipid membrane dye (red) after aldosterone treatment.

We used aldosterone (see figure legends where applicable) to stimulate sodium transport and increase the likelihood of imaging transport phenomena. Aldosterone increases surface abundance of both ENaC and Na-K ATPase, by preventing recycling of the channels (1).

To investigate the vesicles in more detail, cells were co-stained with additional cell markers. Figure 1C shows the co-staining of live mCCD_cl1_ cells with NaG and tubulin dye (light blue). The cell, situated on the side of a dome, contains NaG-rich vesicles framed by cytoskeleton microtubules, suggesting a defined intracellular structure. Cells were then co-stained with NaG and membrane dye (Fig. 1D, red), revealing that NaG-stained vesicles are surrounded by membranes, which further confirms the existence of intracellular structures containing sodium.

We attempted to identify the vesicles by a process of elimination. Cells were first co-stained with NaG and Lysotracker Red, which identifies lysosomes (Fig. 2A, arrow heads). Many small intracellular structures co-stained for NaG and lysotracker, as lysosomes are known to contain Na^+^ (7). We then co-stained with NaG and Mitotracker Red, which identifies mitochondria. No co-staining could be seen for mitochondria. More generally, the structures stained with NaG ranged in size from 0.4 to 6.4μm (see size distribution in Fig. 2B), while lysosomal staining showed a consistent diameter of ∼0.5μm. We can reasonably conclude that the bigger vesicles stained with NaG are neither lysosomes nor mitochondria.

**Figure 2.**
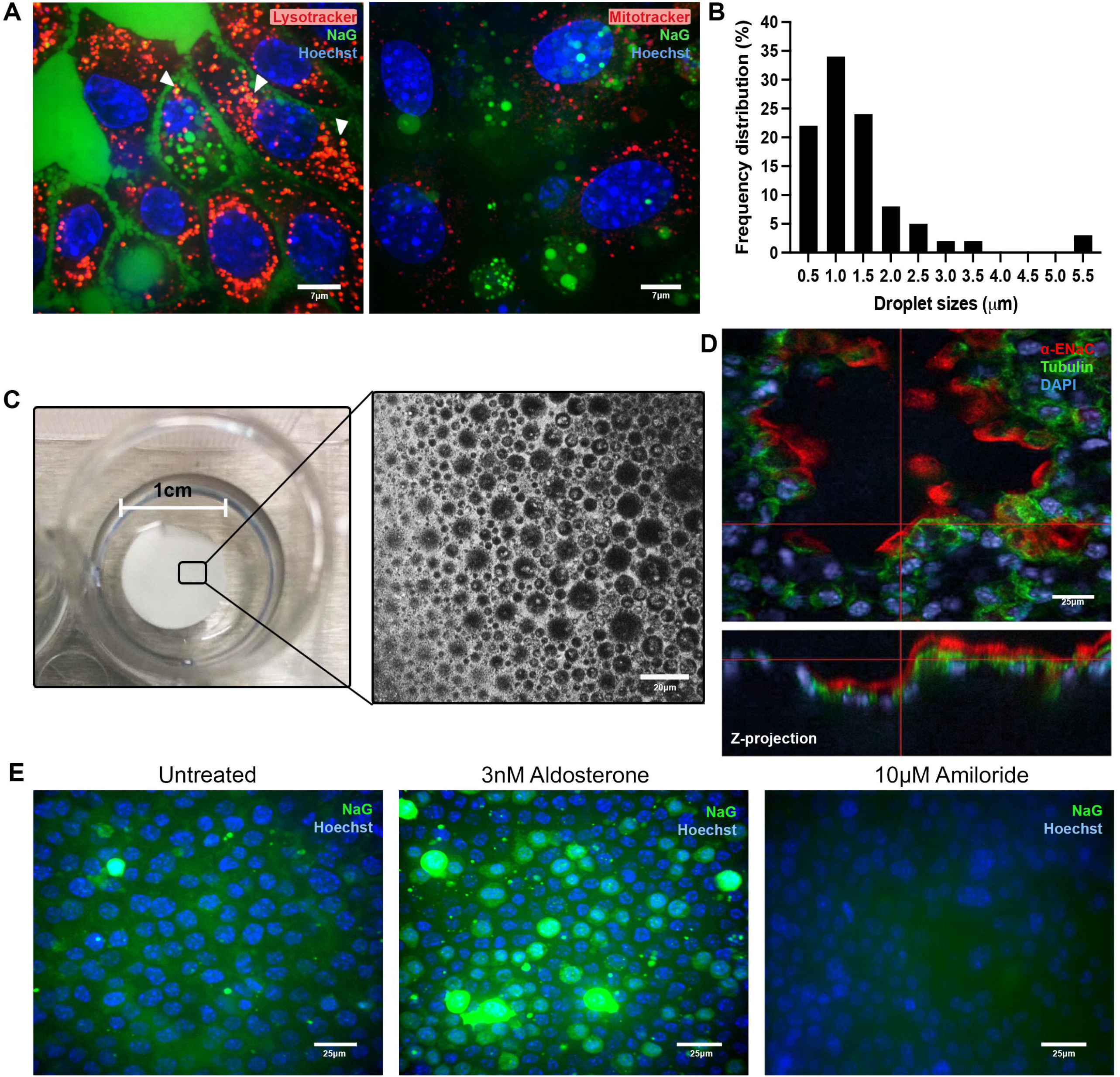
A. Left, staining of live mCCD_cl1_ cells using NaG (green) and Lysotracker (red) after aldosterone treatment. Cell nuclei in blue (Hoechst). Arrowheads show examples of co-staining (small yellow dots). Right, staining of live mCCD_cl1_ cells using NaG (green) and Mitotracker (red). B. Size distribution in percentage of NaG^+^ structure diameters (measured in mCCD_cl1_ cells treated with 3nM Aldosterone for 3 hours). C. Image of 3D polyHIPE scaffold shown in the well of a cell culture dish. Right, magnified view of the surface of the scaffold imaged using the substrate auto-fluorescence. D. Immunostaining of fixed mCCD_cl1_ cells cultured on 3D scaffold using an anti-αENaC antibody (red) and anti-acetylated-αTubulin antibody (green). Z-projection shows the roughness of the scaffold surface and the apical localisation of αENaC staining. Cell nuclei in blue (DAPI). E. Staining of live mCCD_cl1_ cells cultured on 3D scaffold using NaG. Cell were imaged untreated (left), treated with 3nM Aldosterone for 3h (middle) or 10μM Amiloride for 10min (right). Cell nuclei in blue (Hoechst).

To observe potential intracellular sodium transport dynamics, we cultured mCCD_cl1_ cells on bespoke polyHIPE scaffolds, which allow for polarization and transcellular transport, and offer a more convenient solution for handling and imaging than filters such as Transwells (see Figs. 2C and 2D). Moreover, recent advances in “organ-on-a-chip” technologies have shown that cells cultured in a 3D environment have an increased survival and tend to be more differentiated than the same cell lines cultured in classic 2D settings (8). This is particularly relevant in the kidney, where physiological mechanisms are tightly linked to the complex 3D organisation of renal structures, and to cell polarisation.

When treated with aldosterone on this substrate, we observed a “recruitment” of mCCD_cl1_ cells taking up sodium (see Fig. 2E), which is consistent with a rapid translocation of existing ENaC to the membrane and activation of sodium transport mechanisms. The wide range of fluorescence intensities shown in the cell population is also consistent with the extreme variability in expression of principal cell markers observed previously in mCCD_cl1_ (9), and also in primary collecting duct cells using single cell RNA sequencing (10). NaG staining was absent, as expected, when cells were treated with amiloride, which blocks ENaC, prior to NaG incubation (Fig. 2E).

To further investigate NaG staining, we used an equivalent fluorescent dye for potassium, ‘Potassium green’ (KG), on doming cells. Contrary to NaG, KG was more evenly distributed in the cytoplasm, though notably excluded from intracellular structures of a size equivalent to the Na^+^ vesicles. Extracellular KG intensity was low in the filtrate inside the domes, an observation in keeping with the opposite direction of flux of Na^+^ and K^+^ in these cells (Fig. 3A). When doming cells were treated with ouabain (an Na/K ATPase blocker, 3h at 1uM) before imaging with NaG (Fig. 3B) the ouabain-treated cells show collapsing domes, and NaG stain concentrated in the cells rather than the filtrate. These results further validate the ability of NaG staining to faithfully report principal cell Na^+^ biology.

**Figure 3.**
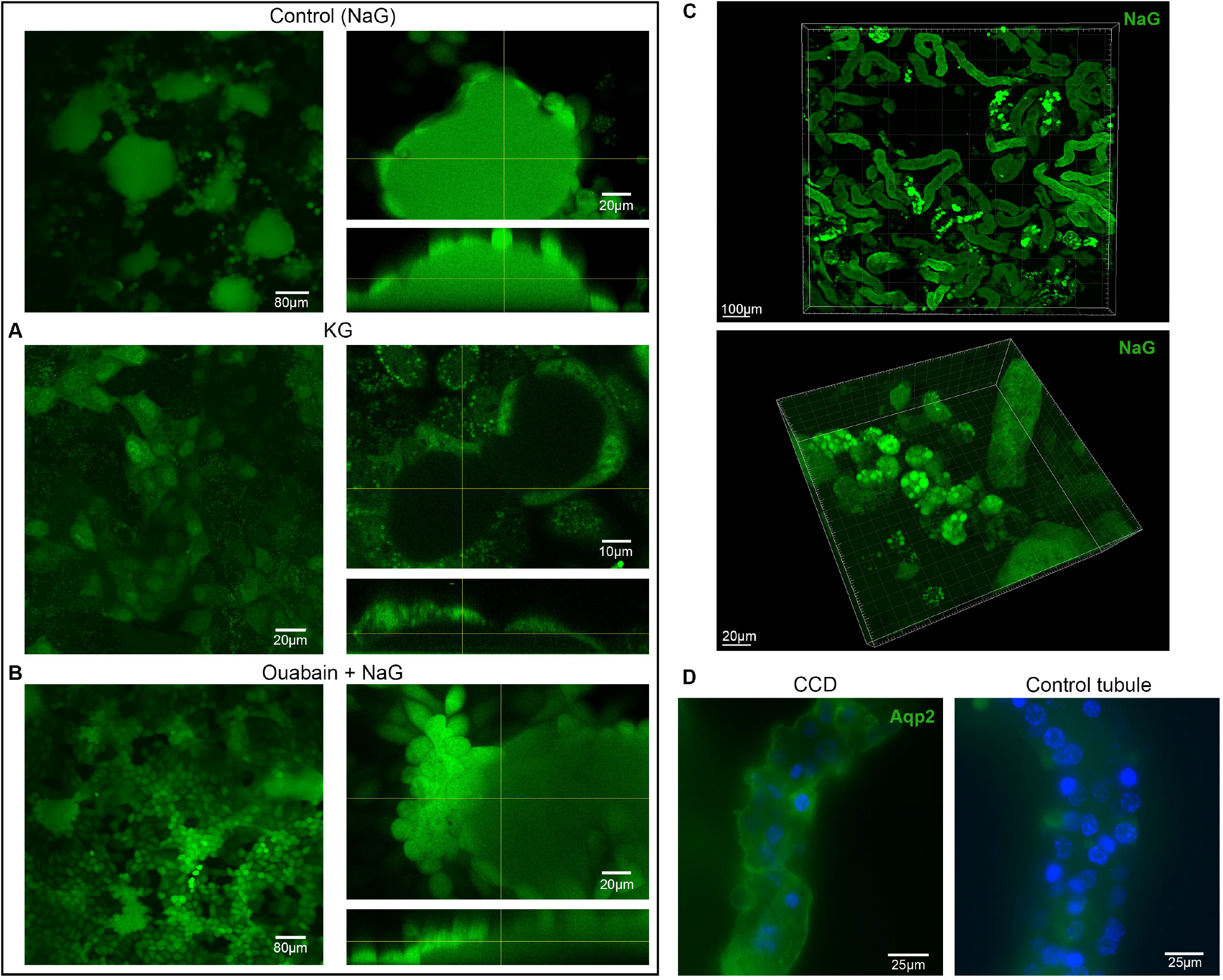
Top left, control figure of doming mCCD_cl1_ cells with NaG for comparison A and B. A. Confocal images (right: with z-projection of a dome) of doming mCCD_cl1_ cells after incubation with KG dye. B. Confocal images of doming mCCD_cl1_ cells after incubation with ouabain and NaG. On the right, a close-up image with z-projection shows one of the collapsed domes. C. 3D reconstruction views (Imaris software) of isolated tubule suspension from mouse kidney after incubation with NaG. High magnification images (bottom) show that the bright punctate staining observed on the top image corresponds to vacuolation inside the cells of some tubules. D. Immunostaining of mouse kidney tubules with anti-Aqp2 antibody (green) shows staining in similar tubule segments as identified in (C). Other tubules showed light to absent staining (right).

To investigate NaG staining *ex vivo*, whole kidney tubules were isolated from a WT mouse and incubated in NaG. CCD were identified by shape (cobblestone-like segment, branching segments), and validated by Aqp2 staining in separate, fixed samples (Fig. 3C-D). Confocal microscopy reveals similar ‘vesicle’ staining in these nephron segments, while other tubule segments displayed more diffuse staining. While our results in primary cells are preliminary, our observations confirm that the Na^+^ pattern seen in mCCD_cl1_, is specific to collecting duct cells.

To observe potential vesicle dynamics, we performed time-lapse imaging on cells cultured on 3D scaffolds described above. Confocal microscopy was used to acquire Z-stack images of cells of interest at high magnification. Image acquisition frequency - every 2 to 4 min - was dependent on sample size, and the number of channels imaged.

Time-lapse movies show a very dynamic environment, with NaG^+^ vesicles both increasing and decreasing in fluorescence intensity. In Fig 4A (and Movie S1), images over time show a 5μm diameter vesicle releasing NaG in a localised manner over the course of ∼20min. Fig 4B (and Movie S2) also shows NaG staining in a vesicle directionally decreasing, suggesting a controlled ‘emptying’ of the vesicle contents. The dynamics of the vesicles contrast with the smaller NaG^+^ structures surrounding them, which are typical, both in size and movements, of lysosomes (11). Fig 4C (and Movie S3) shows the increase of NaG staining in a vesicle (indicated with a blue arrow) and the fading of the staining in another (indicated with a red arrow), suggesting the presence of distinct regulatory mechanisms. Similar dynamics were also observed on doming cells (data not shown). In Figure 4A and B, Hoechst +/- Lysotracker were used on the samples in addition to NaG; only the NaG channel is shown for clarity.

**Figure 4.**
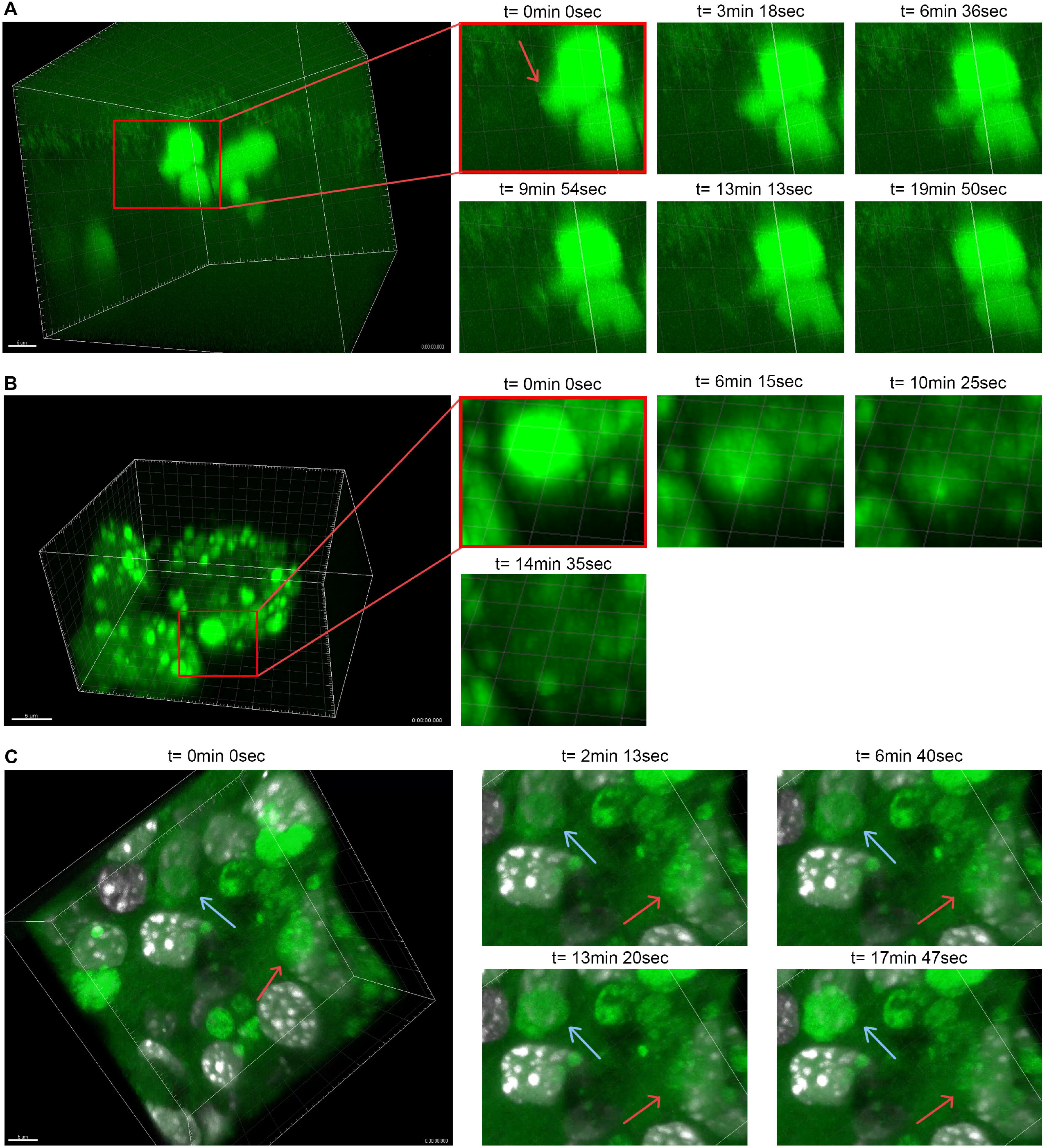
NaG staining in mCCD_cl1_ cells cultured on 3D polymer scaffolds after aldosterone treatment. A. Left panel: 3D reconstruction of confocal images at time t = 0. Right panels: area of interest magnified and shown between t0 and 20 mins (taken from video S1). NaG movement is indicated (red arrow). B. Left panel: 3D reconstruction of confocal images at time t = 0. Right panels: area of interest magnified and shown between t0 and 15 mins (taken from video S2). C. Left panel: 3D reconstruction of confocal images at time t = 0, showing a vesicle with increasing fluorescence (blue arrow) or decreasing fluorescence (red arrow). Right panels: area of interest magnified and shown between t0 and 18 mins (taken from video S3).

## Discussion

In this study we provide evidence for a novel, regulated and dynamic compartmentalisation of sodium in CCD cells. These observations will seed further detailed mechanistic investigations to understand how sodium transits the cytoplasm in Principal cells of the kidney collecting duct.

Corona Green AM has previously been used as a sodium indicator in mammalian, zebrafish, and plant cells (12–14). Of particular note is a recent study of electrolytes in mouse embryos, where sodium ions were shown to accumulate in the blastocoel cavity over a 40 minute incubation period (15). We believe that vesicles have not been reported in other mammalian cells to date. However, the compartmentalisation of sodium into vacuoles, to avoid toxic concentrations in the cytosol is a wellknown phenomenon in plants (14), which also regulate Na^+^ H-ATPase activity at the cell membrane according to sodium concentration (16). Since we saw evidence of vesicles in CD cells *ex vivo*, we therefore do not believe that the vesicles seen in the mCCD cells per se are an artefact of the dye treatment, but further studies are needed to eliminate doubts. More specifically, potential experiments include using alternative fluorescent indicators (though as indicated before, SBFI showed limited uptake), dropping incubation temperature, and using a range of NaG concentrations. It would also be very valuable to conduct *in vivo* studies, such as injection of NaG in mice and consequent tissue harvest in control *versus* increased sodium uptake.

There are a number of caveats to this study. Plasticity of the cortical collecting duct between PC and IC cells, together with the presence of transitional cells, has been demonstrated *in vivo* (17). Since mCCD cells exhibit plasticity in phenotype, expressing a spectrum of both PC and IC markers, with recruitment of PC markers to the luminal and apical membranes following aldosterone treatment, they were used in the present study. NaG is not a ratiometric dye, and is only visible when sodium concentration exceeds ∼10mM according to the manufacturer, with a Kd previously measured at 80mM (12). Though ratiometric dyes exist (such as SBFI), they exhibited very limited uptake by the mCCD_cl1_ cells, which may be due to a higher molecular weight. The main caveat is the potential effect of NaG itself on the cells. It is possible that the dye may accumulate in intracellular compartments due to the dye concentration and incubation temperature, or the high sodium transport characteristic of these cells. It is also possible that aldosterone affects NaG cleavage, accumulation, and fluorescence. However, it should be noted that NaG dye accumulates in domes, meaning that NaG is still effectively exported from the mCCD cells.

We considered early on the possibility that the cells are stressed and apoptotic or autophagic. However apoptotic cells present a different phenotype (18), and neither autophagic bodies nor stress granules (which are biomolecular condensates lacking a membrane)(19) were observed in previous studies involving these cells.

In addition to the obvious discrepancy in size range, we applied a process of elimination to conclude that the vesicles are neither lysosomes nor mitochondria but are surrounded by lipid membranes and bordered by microtubules. Most vesicles were situated towards the basolateral membrane. Cytoplasmic Na^+^ concentration may be insufficient to cause reporter fluorescence, implying that Na^+^ is concentrated in the vesicles. Evidence suggests that the Na^+^ vesicles exclude K^+^, since the size of the observed K ‘negative’ volumes corresponded to the size of the Na^+^ vesicles. It is probable that these structures are Na^+^-specific, but further studies are necessary.

We can speculate that Na^+^ is transported through ENaC, then through cargo transport (20) or ionic currents (21) along the cytoskeleton and deposited/concentrated in vesicles before being transported across the Na-K ATPase channel. This suggests regulatory mechanisms controlling Na^+^ deposition into and release from the vesicles. It has been shown that aldosterone, stimulates sodium transport across the PC, by inhibiting p38 kinase-mediated endocytosis of Na-K ATPase (22). We saw a significant increase in both the number and size of vesicles following aldosterone treatment.

Confocal imaging in Figure 4 indicates that the vesicles have internal structure. This would lead us to speculate that they may be multivesicular bodies, which are a specialised subset of endosomes that can contain both membrane-bound intraluminal vesicles and exosomes (23). Multivesicular bodies either pass their contents to lysozymes for subsequent degradation, or to the plasma membrane for expulsion of exosomes. Though we were limited to imaging every 2 to 4 min, meaning that rapid movements would not be detected, we may have captured each of these events in movies S1 and S2, respectively. By analogy, the inclusion of intraluminal vesicles into the multivesicular body may have been captured in movie S3. Evidence of multivesicular bodies in CD sections has been reported previously (1).

The presence of NaG structures in primary CCD cells show that our findings are not unique to mCCD cells and are relevant to native principal cells. Preliminary observations using electron microscopy did not yield further insights, however the study was limited. Given their transient nature and the fact that not all mCCD cells exhibit the phenomenon (consistent with their known plasticity), an extensive electron microscopy screen in the presence and absence of aldosterone and NaG, may be informative. The possibility that these sodium-containing subcellular structures appear as a consequence of NaG itself must be considered, however we do not believe that the presence of vesicles per se is an artefact of NaG because such vesicles have not been observed in other cells using this stain. It is entirely possible that NaG delays, or affects, the filling and emptying of the vesicles allowing us to visualise them. Our data suggest the presence of sodium-containing vesicles within principal cells and that NaG may be an informative tool to investigate the dynamics of sodium transport across the principal cell, and, possibly, other sodium transporting cells.

In conclusion, we have demonstrated a potentially novel intracellular compartment in sodium-transporting mCCD_cl1_ cells, with wide implications for regulation of intracellular sodium handling in the CCD, a key nephron segment in blood pressure regulation. While further work is needed to confirm the results both in primary cells, and in vivo, these observations open a new field of study for the regulation of sodium compartmentalisation in the collecting duct and offer important new insights into intracellular sodium dynamics.

## Acknowledgments

We acknowledge Professors Chris Gregory, Margarete Heck, David Lyons, Mike Shipston and Sebastian Bachmann, and Dr Bryan Conway for helpful discussions and Lynsey Melville and Dr Steve Mitchell for assistance with transmission electron microscopy.

## Supplemental data

Supplemental movies are available by following the private links:

Movie S1: https://figshare.com/s/722c08e39948336b7859

Movie S2: https://figshare.com/s/d0be7a7c6a623abf18ca

Movie S3: https://figshare.com/s/12731a3547b4931e5385

